# Simulated microgravity alters short-term evolutionary trajectories of Orsay virus in *Caenorhabdidits elegans*

**DOI:** 10.64898/2026.05.14.725097

**Authors:** Ana Villena-Giménez, Victoria G. Castiglioni, Santiago F. Elena

## Abstract

**Background:** Environmental conditions shape the evolutionary trajectories of RNA viruses, yet little is known about how complex physical stressors such as microgravity influence host-virus interactions and viral evolution. Here, we investigated the short-term evolutionary consequences of simulated microgravity on the *Caenorhabditis elegans* - Orsay virus (OrV) system.

**Methods:** OrV was subjected to six serial passages in hosts acclimated to low-shear modeled microgravity, with parallel evolution under standard-gravity. Evolutionary outcomes were evaluated using virulence, transmission, and replication traits, all measured under standard-gravity conditions.

**Results:** Viral load fluctuated across passages in both environments, with lower mean accumulation in microgravity-evolved lineages. After evolution, we detected no significant changes in virulence. Transmission increased in standard-gravity lineages but not in microgravity-evolved ones, while viral replication decreased in all lineages, with a stronger decline in those evolved under microgravity. However, the magnitude of phenotypic changes was generally modest.

**Discussion:** These results indicate that evolution under microgravity can alter viral phenotypic trajectories over short timescales. However, because all traits were assayed under standard-gravity conditions, we cannot directly assess local adaptation to microgravity, and the observed differences may reflect environment-specific trade-offs rather than reduced fitness *per se*. Furthermore, the limited number of passages and the modest magnitude of phenotypic change suggest that evolutionary responses may still be in an early stage.

**Conclusion:** Overall, our findings provide initial evidence that simulated microgravity can influence the evolutionary dynamics of an RNA virus, while highlighting the need for reciprocal fitness assays and longer-term experiments to fully characterize adaptation to altered gravitational environments.

## 1 Introduction

Human activities in space are rapidly expanding, with renewed interest in long-duration missions, lunar exploration, and the eventual establishment of permanent settlements beyond Earth. These scenarios raise important biological and medical questions, because organisms exposed to space-like environments experience combinations of physical stressors that differ substantially from those on Earth. Among these, reduced gravity is one of the most pervasive and biologically relevant factors. Microgravity has been associated with alterations in multiple physiological systems, including brain structure, cardiovascular function, muscle maintenance, and immune regulation (Stein 2013; Buchheim et al. 2019; Roberts et al. 2019; Azariah and Terranova 2025). Understanding how microgravity affects host physiology is therefore essential for anticipating health risks during spaceflight.

One particularly relevant consequence of spaceflight-associated physiological stress is altered host-pathogen interaction. Immune dysfunction during spaceflight has been proposed to contribute to the reactivation of latent viruses in astronauts, including varicella-zoster virus, cytomegalovirus and Epstein-Barr virus (Cohrs et al. 2008; Mehta et al. 2014, 2017). Although these observations indicate that spaceflight can affect viral infection outcomes, the mechanisms involved remain incompletely understood. In particular, it is still unclear whether reduced gravity influences viral infections mainly through changes in host physiology, through direct effects on viral particles or replication processes, or through a combination of both. Addressing this question is important not only for astronaut health, but also for understanding how novel physical environments may reshape host-virus interactions.

Experimental systems that simulate microgravity provide a tractable way to disentangle some of these effects under controlled laboratory conditions. Devices such as random positioning machines (RPM) allow organisms to be exposed to low-shear modeled microgravity, making it possible to examine how altered gravitational cues affect development, physiology, immunity, and infection. The nematode *Caenorhabditis elegans* is especially useful in this context because it is experimentally tractable, genetically well characterized, and has been widely used to study stress responses, immunity, and host-pathogen interactions. Importantly, several effects of spaceflight or simulated microgravity observed in *C. elegans* resemble broader responses seen in animals, including changes in immune and metabolic pathways (Morukov et al. 2011; Garrett-Bakelman et al. 2019; Çelen et al. 2023; He et al. 2024).

The natural pathosystem formed by *C. elegans* and Orsay virus (OrV) provides a particularly valuable model for studying viral infection and evolution under altered environmental conditions. OrV is a natural RNA virus of *C. elegans*, and previous studies have characterized several aspects of its interaction with the host, including infection dynamics, host transcriptional responses, and disease-related traits. Recent work has shown that simulated microgravity affects both *C. elegans* life-history traits and OrV infection outcomes, including changes in viral accumulation and host responses (Villena-Giménez et al. 2026a, 2026b). These findings indicate that reduced gravity can modify the ecological context in which OrV replicates and transmits, raising the possibility that it may also influence viral evolutionary dynamics.

RNA viruses are especially appropriate for addressing this question because their large population sizes, short generation times, and high evolutionary rates allow rapid responses to environmental change (Domingo et al. 1996; Elena and Sanjuán 2007; Elena et al. 2008). Experimental evolution studies have repeatedly shown that viral populations can adapt to novel or stressful environments, although such adaptation often involves trade-offs among replication, stability, transmission, and virulence. For example, thermal stress, altered host proteostasis, and other environmental pressures can reshape viral fitness landscapes and favor different genetic or phenotypic solutions (Phillips et al. 2017; Singhal et al. 2017). These observations suggest that microgravity, by altering either the host environment or the physical conditions experienced by the virus, could modify the selective pressures acting on viral populations.

Despite this relevance, little is known about how microgravity affects the evolution of RNA viruses. Most available studies have focused on the immediate physiological or infection-related consequences of spaceflight or simulated microgravity, rather than on how viral populations change across successive generations in such environments. This represents an important gap, because infection outcomes observed at a single time point may differ from those produced after viral populations have had the opportunity to evolve. Even short-term evolution may alter key viral traits such as replication, transmission or virulence, and may reveal whether reduced gravity imposes novel constraints or redirects viral evolutionary trajectories.

Here, we performed an evolution experiment to test the effect of microgravity, simulated employing a random positioning machine, on the short-term evolution of OrV in its host *C. elegans*. OrV was serially passaged for six rounds in animals maintained either under standard-gravity or simulated microgravity conditions. At each passage, fresh animals were inoculated with viral populations collected from the previous passage, and viral replication proceeded for 44 hours post-inoculation (hpi), corresponding to larval development of the host. After the sixth passage, the five independent lineages evolved under standard-gravity and microgravity conditions were evaluated in terms of virulence, transmission and replication rate under standard-gravity assay conditions.

## 2 Materials and Methods

### 2.1 *C. elegans* strains and culturing

*C. elegans* were cultured and maintained at 20 °C on nematode growth medium (NGM) agar plates seeded with *E. coli* OP50. The transgenic strain ERT54 (*jyIs8[pals-5p::GFP + myo-2p::mCherry]X*), which carries a wildtype Bristol N2 genetic background and expresses GFP in response to intracellular infection (Bakowski et al. 2014), was used for all experiments. The more susceptible strain SFE2 (*drh-1(ok3495)IV; mjIs228*) was used to generate OrV stocks.

To synchronize populations, plates containing embryos were gently washed with M9 buffer to remove larvae and adults while leaving the embryos behind. After 1 h, plates were washed again with M9 buffer to collect larvae that had hatched during that interval, and these were transferred to freshly seeded NGM plates.

### 2.2 Microgravity simulation

An RPM (Yuri Gravity GmbH, Meckenbeuren, Germany) was used to simulate microgravity conditions by operating in zero-gravity mode. G-force values were monitored using the RPM software and maintained at approximately 0.001 - 0.002 g throughout the experiments. The RPM was housed inside an incubator set to 20 °C.

Animals used in microgravity experiments were acclimated for two generations, and all assays were performed on the third generation and its progeny. Plates were sealed with Parafilm to maintain humidity.

### 2.3 Experimental evolution of OrV

Fig. 1 presents a schematic overview of the experimental evolution design. Four hundred L1 larvae were inoculated with 60 µL of JUv1580_vlc and collected 44 hpi. If fewer than 50% of the animals expressed the *pals-5p::GFP* reporter, the passage was discarded and a new population was inoculated using the viral stock from the previous successful passage. Conversely, when more than 50% of animals showed *pals-5p::GFP* expression, nematodes were collected in 1.8 mL of M9 buffer, incubated for 10 min at room temperature, vortexed, and centrifuged for 2 min at 400 g. The nematode pellet was stored at −80 °C, while the supernatant was centrifuged twice at 21,000 g for 5 minutes at 4 °C and then filtered through a 0.2 µm membrane.

**FIGURE 1.**
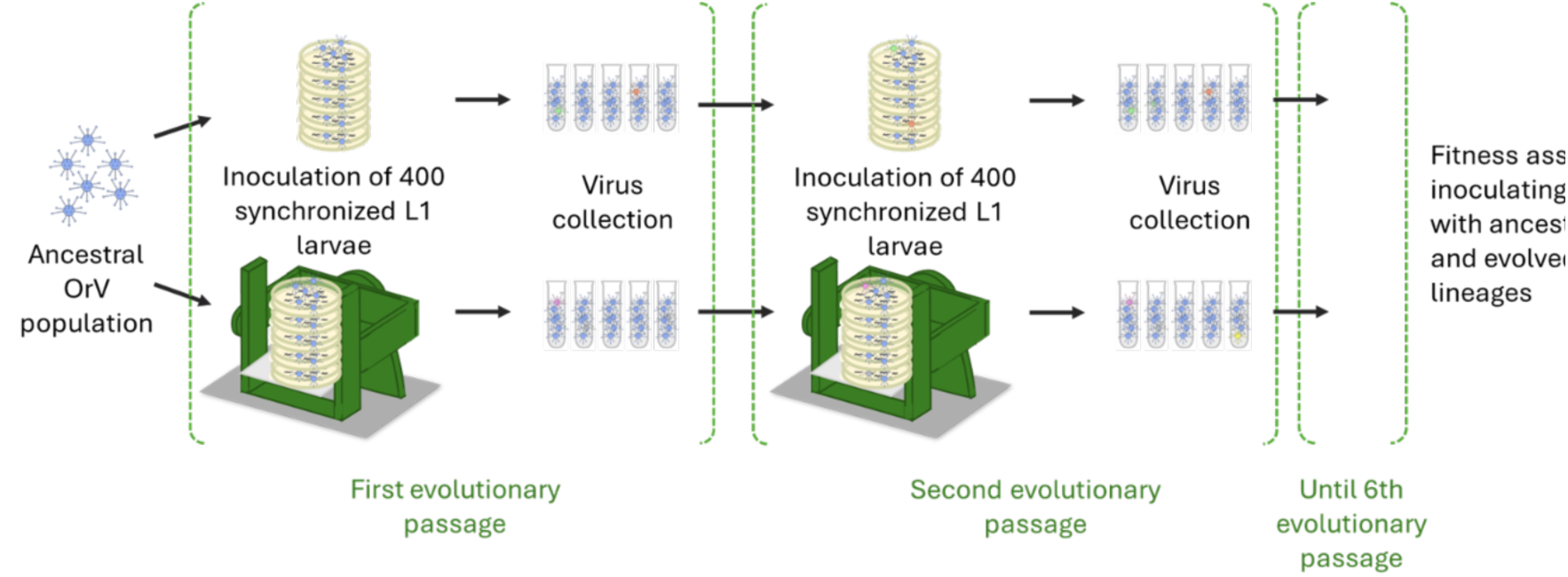
Evolution under standard-gravity and simulated microgravity conditions was conducted in parallel. For the microgravity treatment, viral evolution was performed in third-generation animals acclimated to simulated microgravity, in order to reduce the contribution of acute host stress responses to viral evolutionary dynamics.

A total of 60 µL of the resulting viral preparation was used to inoculate fresh L1 larvae, and this process was repeated for six passages in five independent evolution lineages. After the 6^th^ passage, viral concentration in the supernatant was quantified as described below. RNA from the final viral stock was extracted using the Viral RNA Isolation Kit (NZYTech, Lisboa, Portugal). Viral RNA concentration was then determined by RT-qPCR using a standard curve. Primer sequences for these RT-qPCR assays are available elsewhere (Castiglioni et al. 2024, 2025; Villena-Giménez et al. 2026a).

### 2.4 Viral stock preparation, virus quantification and inoculation procedure

For preparation of OrV (isolate JUv1580_vlc) stocks, SFE2 animals were inoculated as previously described (Castiglioni et al. 2024). Briefly, animals were allowed to grow for 5 d and then resuspended in M9 buffer (0.22 M KH_2_PO_4_, 0.42 M Na_2_HPO_4_, 0.85 M NaCl, 1 mM MgSO_4_). The suspension was allowed to stand for 15 min at room temperature, vortexed, and centrifuged for 2 min at 400 g. The supernatant was then centrifuged twice at 21,000 g for 5 min and passed through a 0.2 µm filter. Viral RNA from the resulting stock was extracted using the Viral RNA Isolation Kit (NZYTech). Viral RNA concentration was measured by RT-qPCR using a standard curve and normalized across batches (details below).

To generate the standard curve, cDNA of JUv1580_vlc was synthesized using AccuScript High-Fidelity Reverse Transcriptase (Agilent, Santa Clara CA, USA) and reverse primers annealing near the 3’ end of the genome. Approximately 1,000 nucleotides from the 3’ end of RNA2 were amplified using forward primers containing a 20 nucleotides T7 promoter sequence and DreamTaq DNA Polymerase (Thermo Fisher Scientific, Waltham MA, USA). PCR products were gel-purified using the MSB Spin PCRapace kit (Invitek Molecular GmbH, Berlin, Germany), followed by *in vitro* transcription with T7 RNA Polymerase (Merck, Rahway NJ, USA). Remaining template DNA was digested with DNase I (Thermo Fisher Scientific). RNA concentration was measured using a NanoDrop spectrophotometer (Thermo Fisher Scientific), and RNA copy number per µL was calculated using the EndMemo RNA Copy Number Calculator (https://www.endmemo.com/bio/dnacopynum.php). Primer sequences used for generating the standard curve are reported in Castiglioni et al. (2024).

Inoculation of the first evolutionary passage was performed as follows. Synchronized populations were infected by pipetting 60 µL of viral stock directly onto the bacterial lawn containing the animals. The normalized inoculum contained 2.6×10^7^ copies of OrV RNA2 per µL. The efficiency of this viral stock, measured as the proportion of animals showing activation of the *pals-5p::GFP* reporter at 48 hpi, was 72 ±3% (mean ±SEM; *n* = 5 plates, 44 - 48 animals per plate).

Pellets of animals collected at 44 hpi throughout the experimental evolution were stored at −80 °C. RNA extractions were performed as described above. However, viral load quantification by RT-qPCR used a separate standard curve-based determination prepared as noted previously; the procedure for generating and applying this standard curve is described in detail above.

To determine the concentration of inocula used for downstream experiments, supernatants from the final passage of each lineage were processed as described above. Specifically, RNA was extracted using the Viral RNA Isolation Kit (NZYTech) and quantified by RT-qPCR using a standard curve. Lineages with concentrations greater than 10^5^ copies of RNA2 per µL were diluted to this concentration in a final inoculation volume of 60 µL. Conversely, if a lineage contained fewer than 10^5^ copies/µL, the inoculum volume was increased to reach a total of 6×10^6^ RNA2 copies/µL.

### 2.5 RNA extractions and RT-qPCRs

A total of 500 µL of TRIzol (Thermo Fisher Scientific) was added to the frozen worm pellet, which was then disrupted by five cycles of freeze-thawing followed by five cycles of 30 s of vortexing and 30 s of rest. Next, 100 µL of chloroform was added, and the tubes were shaken for 15 s and left to stand for 2 min. Samples were centrifuged for 15 min at 11,000 g at 4 °C, and the upper aqueous phase containing RNA was mixed with an equal volume of 100% ethanol. The mixture was loaded onto RNA Clean and Concentrator columns (Zymo Research, Irvine CA, USA), and the remaining steps were carried out according to the manufacturer’s instructions.

RT-qPCR assays were performed using qPCRBIO SyGreen 1-Step Go Hi-ROX (PCR Biosystems, London, UK) on an ABI StepOne Plus Real-Time PCR System (Thermo Fisher Scientific). A total of 10 ng of RNA was used per reaction.

### 2.6 Viral load from evolved lineages

Five replicates of synchronized populations of 500 inoculated animals were collected at 14 hpi using PBS containing 0.05% Tween. All evolved lineages were inoculated under standard-gravity conditions, and each inoculum contained an estimated 6×10^6^ copies of OrV RNA2, as determined by RT-qPCR. Samples were centrifuged for 2 min at 1,350 rpm, and the supernatant was discarded. Two additional wash steps were performed before the samples were flash frozen in liquid N_2_. RNA extractions were carried out as described in Section 2.3, followed by RT-qPCR analysis.

### 2.7 Progeny assay

L4 larvae were individually picked from a synchronized population and transferred to separate fresh plates. Animals were moved to new plates every 48 h until egg-laying ceased. Progeny were counted 48 h after the adult had been removed from the plate. All assays were performed under standard-gravity conditions.

For OrV-infected conditions, synchronized L1 larvae were inoculated with 6×10^6^ copies of RNA2, and infected L4 larvae were specifically selected using the *pals-5p::GFP* fluorescent reporter. GFP signal was assessed using a Leica MZ10 F fluorescence stereomicroscope equipped with a Flexacam C3 camera, a 10×/23B objective, and mCherry M10F/MZ FLII and GFP3 MZ10F filter sets (Leica Microsystems GmbH, Wetzlar, Germany).

### 2.8 Transmission assay

A variable number of L1 larvae were inoculated with 6×10^6^ RNA2 copies. After 24 h, 20 infected animals exhibiting *pals-5p::GFP* fluorescence were transferred to a plate containing 50 L1 larvae. Twenty-four h after transfer, the 20 infected animals were removed, and the remaining worms were scored as infected or non-infected based on the presence or absence of GFP signal, respectively. These counts were used to calculate the percentage of infected animals. The experiment was performed in five replicates under standard-gravity conditions.

GFP fluorescence was assessed using a Leica MZ10 F fluorescence stereomicroscope equipped with a Flexacam C3 camera, a 10×/23B objective, and mCherry M10F/MZ FLII and GFP3 MZ10F filter sets (Leica Microsystems).

### 2.9 Statistical analysis

Viral load was analyzed using a generalized linear model (GLM) with a Gamma distribution and a log-link function. Experimental treatment and evolutionary passage were included as orthogonal fixed factors, and independent evolutionary lineages were treated as a random factor nested within experimental treatment.

Progeny from animals infected with ancestral and evolved OrV lineages was analyzed using a GLM with a negative binomial distribution and a log-link function. Experimental treatment was included as a fixed factor, and evolutionary lineages were included as a random factor nested within the fixed factor.

Transmission efficiency data were analyzed using a GLM with a binomial distribution and a logit-link function, again with experimental treatment as a fixed factor and evolutionary lineages as a random factor nested within it.

In all cases, pairwise differences were assessed using the post hoc sequential Bonferroni method. All three sets of analyses were performed using SPSS version 30.0.0.0 (IBM Corporation, Armonk NY, USA).

To estimate rates of phenotypic evolution for each trait, individual time-series data were modeled using ARIMA(*p*, *d*, *q*) following the procedures described in Elena et al. (2025). Model selection was performed using the R package forecast version 8.23.0, including automatic estimation of the Box-Cox transformation parameter λ. Overall trends in time-series data were estimated using the robust Sen-Theil nonparametric slope estimator as implemented in the R package trend version 1.1.6. Slope significance was evaluated using the Mann-Kendall 1″ test with the R package Kendall version 2.2.2. Differences in evolutionary rates between standard and microgravity conditions were evaluated using a two-sample *t-*test assuming unequal variances.

The following quantitative genetics parameters were estimated using custom R scripts for each trait and evolutionary condition: additive genetic variance (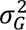), broad-sense heritability (*h*^2^), selection gradient (*S*), response to selection (*R* = *Sh*^2^), and the effective minimum number of loci contributing to each trait (*n_min_*). Error estimates were obtained using 10,000 bootstrap pseudosamples. Differences between treatments for each parameter were assessed using two-sample *t*-tests.

R coding was done using R version 4.5.3 in RStudio version 2026.04.0+526 (Posit PBC, Boston MA, USA).

## 3 Results

To evaluate phenotypic differences among evolved viral lineages under a common assay environment, we measured three traits associated with OrV performance and infection outcome: viral replication, transmission efficiency and host reproductive output. These endpoint assays were conducted under standard-gravity conditions for both standard-gravity- and microgravity-evolved lineages, allowing direct comparison among evolved populations while avoiding environmental variation during trait measurement. However, this design does not directly test performance in the microgravity environment in which some lineages evolved.

### 3.1 Viral load dynamics during serial passage differ between standard-gravity and microgravity environments

During the experimental evolution, OrV viral load varied substantially across passages in both standard-gravity and simulated microgravity conditions (Fig. 2). Despite these fluctuations, viral accumulation differed consistently between environments. When viral load was analyzed across all passages using a GLM, mean viral load was significantly higher in lineages evolved under standard-gravity than in those evolved under microgravity [(3.3 ±0.3)×10^6^ *vs*. (2.0 ±0.2)×10^6^ RNA2 copies, respectively; χ² = 1122.700, 1 d.f., *P* < 0.001]. Viral load also varied significantly across passagesand the magnitude of this variation depended on evolutionary treatment (χ² = 979.007, 40 d.f., *P* < 0.001). These results indicate that the two environments differed in their overall capacity to support OrV accumulation during serial passage.

**FIGURE 2.**
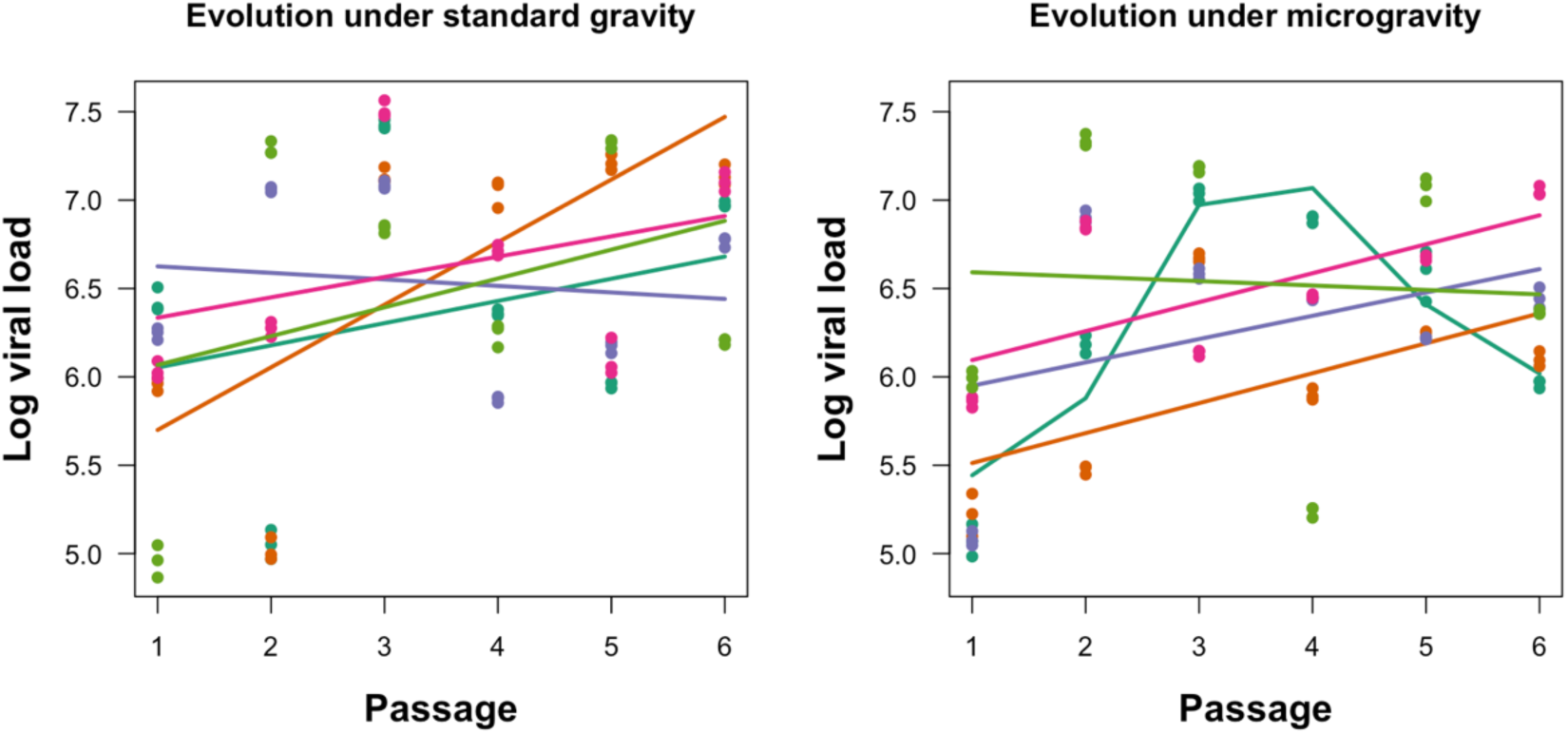
Viral load dynamics across evolutionary passages under standard-gravity (left) and microgravity (right). Dots and line colors distinguish the five independent lineages within each condition. Lines represent the best ARIMA fittings.

To further characterize the temporal structure of viral-load trajectories, we fitted ARIMA models independently to each evolutionary lineage. The best-supported model was ARIMA(1,0,0): *V*_t_ = *C* + *ϕV*_t-1_ + *ɛ*_t_ (Supplementary Table S1), indicating that viral load at a given passage was partly dependent on viral load in the preceding passage. This first-order autoregressive structure suggests that changes in viral accumulation were not purely random from one passage to the next, but instead retained some short-term temporal dependence. Model diagnostics generally supported this approximation, with most lineages showing nonsignificant Ljung–Box tests (*P* ≥ 0.077), indicating limited residual autocorrelation after model fitting.

The autoregressive parameter *ϕ* varied among lineages, suggesting differences in the persistence of viral-load fluctuations. Under standard-gravity, estimates ranged from 0.452 to 0.644, consistent with moderate temporal dependence. Under microgravity, estimates ranged more widely, from 0.478 to 0.976. Notably, one microgravity lineage showed a value close to one, indicating that perturbations in viral load decayed slowly in that lineage. This broader range suggests that microgravity may increase heterogeneity among replicate evolutionary trajectories, although this pattern should be interpreted cautiously given the small number of lineages.

We then estimated temporal trends in viral load using the Sen-Theil slope for each lineage. Most lineage-specific slopes were positive but small, and only one lineage in each treatment showed a significant positive trend: lineage 2 under standard-gravity (Mann-Kendall’s test: *P* = 0.028) and lineage 4 under microgravity (*P* = 0.010). Overall, slope estimates did not differ significantly between standard-gravity and microgravity treatments (*t* = 0.025, 8 d.f., *P* = 0.659). Thus, although viral load differed markedly in magnitude between environments, there was no evidence that the average rate of temporal change in viral load differed significantly between treatments over the six passages.

Finally, viral RNA measured in the supernatants collected after the final passage ranged from 5.8×10^4^ to 1.5×10^6^ RNA2 copies. These final viral stocks were subsequently normalized for the endpoint phenotypic assays described below. Together, these results show that OrV experienced distinct amplification environments during serial passage: viral accumulation was lower under simulated microgravity, and replicate lineages displayed substantial heterogeneity in their temporal dynamics. However, because temporal trends were weak and not significantly different between treatments, these data are best interpreted as evidence that microgravity altered short-term viral-load dynamics, rather than as definitive evidence for directional adaptation during the six-passage experiment.

### 3.2. Replication capacity declines after experimental evolution, especially in microgravity-evolved lineages

After characterizing viral-load dynamics during serial passage, we next evaluated whether the evolved lineages differed in their capacity to replicate under a common assay environment. To this end, we measured viral load at 14 hpi in animals inoculated with the ancestral OrV strain or with each of the evolved lineages. This time point corresponds to the peak of OrV replication previously identified for this system (Villena-Giménez et al. 2026a). All assays were performed under standard-gravity conditions, and inocula were normalized to contain the same number of OrV RNA2 copies. Viral load was quantified by RT-qPCR and expressed as RNA2 genome copies per 10 ng of total RNA. Five biological replicates were analyzed per lineage, except for lineage 4 evolved under standard-gravity, for which four replicates were available.

The ancestral virus showed the highest replication level among the assayed populations (Fig. 3). In contrast, both sets of evolved lineages showed reduced viral loads relative to the ancestor. This decrease was observed after evolution under standard-gravity and was even more pronounced in lineages evolved under simulated microgravity. A GLM confirmed significant differences in viral load among evolutionary treatments (χ² = 588.114, 2 d.f., *P* < 0.001) and among lineages nested within treatments (χ² = 833.649, 8 d.f., *P* < 0.001), indicating that replication capacity varied both according to the evolutionary environment and among independently evolved populations.

**FIGURE 3.**
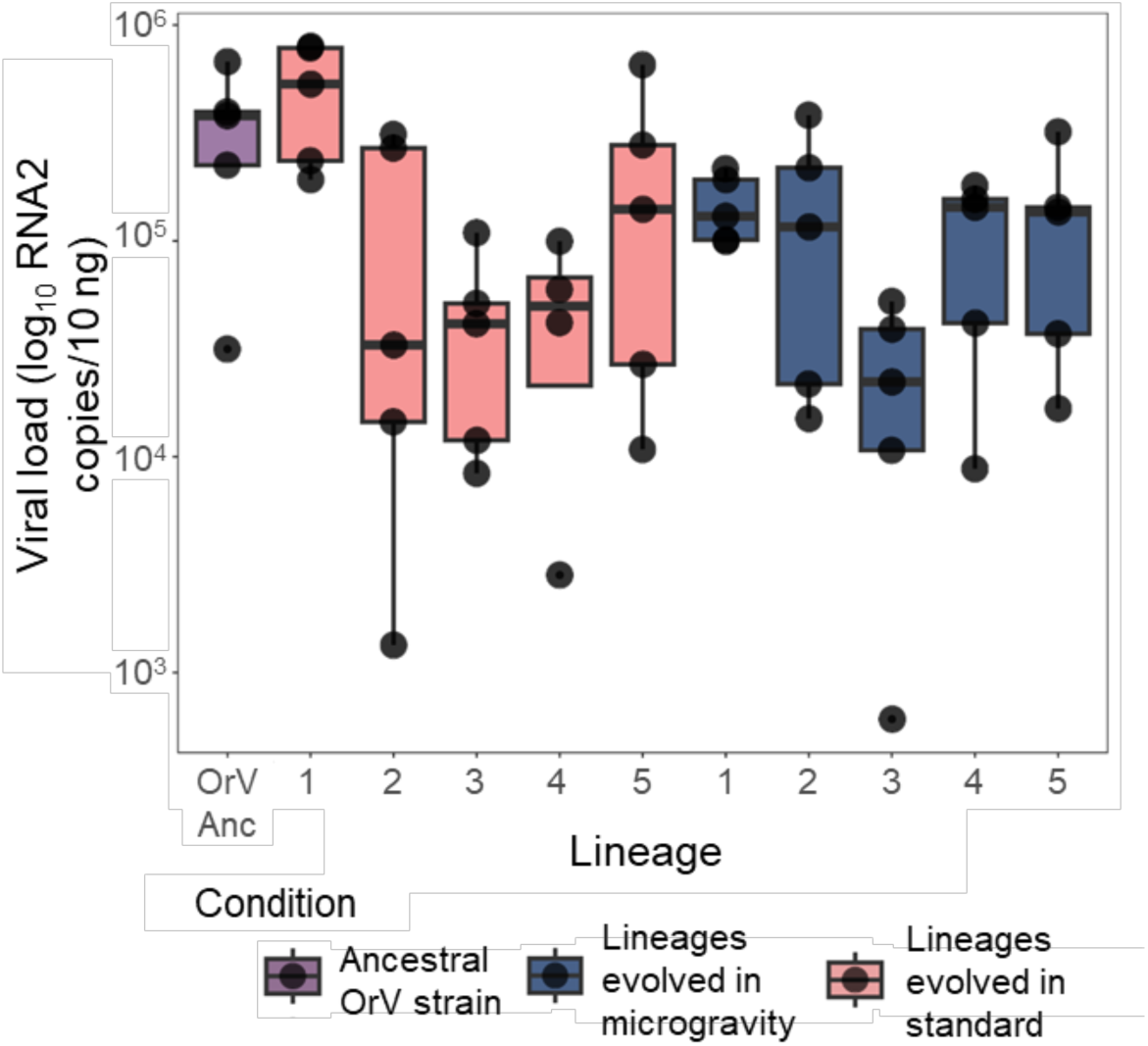
Viral load at 14 hpi of animals inoculated with ancestral OrV (purple) and evolved lineages in standard-gravity (pink) and microgravity conditions (dark blue). Lineages are shown separately, and five replicates were taken per lineage, except for Lineage 4 in standard-gravity conditions in which four replicates were measured.

Pairwise comparisons showed that lineages evolved under standard-gravity had significantly lower viral loads than the ancestral strain [(6.9 ±0.7)×10^4^ *vs*. (2.35 ±0.05)×10^5^ RNA2 copies, respectively; Bonferroni-adjusted *P* < 0.001]. Microgravity-evolved lineages also showed a significant reduction relative to the ancestral strain [(6.00 ±0.06)×10^4^ RNA2 copies; *P* < 0.001]. In addition, viral load differed significantly between the two evolved treatments, with microgravity-evolved lineages exhibiting the lowest replication levels overall (Bonferroni-adjusted *P* < 0.001).

These endpoint measurements are consistent with the lower viral accumulation observed during serial passage under simulated microgravity. However, because replication was assayed only under standard-gravity conditions, the lower viral loads of microgravity-evolved lineages should not be interpreted as direct evidence of reduced fitness in microgravity. Instead, these results indicate that, when evaluated in a common standard-gravity environment, evolved lineages, particularly those previously passaged under microgravity, show reduced replication relative to the ancestral virus. This pattern may reflect trade-offs associated with evolution in the alternative environment, the effects of serial bottlenecks during passage, or limited opportunity for adaptive improvement during the short six-passage experiment.

### 3.3 Transmission increases after evolution under standard-gravity but not after evolution under microgravity

We next examined whether experimental evolution altered the ability of OrV to transmit to new hosts. Transmission efficiency was measured under standard-gravity conditions using animals infected with either the ancestral OrV strain or one of the evolved viral lineages. For each assay, 20 infected donor animals expressing the *pals-5p::GFP* infection reporter were transferred to plates containing 50 uninfected L1 larvae. After 24 h of exposure, donor animals were removed and recipient animals were scored as infected or non-infected based on GFP expression. Transmission efficiency was calculated as the percentage of recipient animals showing the infection reporter. Five biological replicates were performed for each lineage.

Transmission efficiency differed among viral groups (Fig. 4). Lineages evolved under standard-gravity showed higher transmission than the ancestral OrV strain, whereas lineages evolved under simulated microgravity showed intermediate values. A GLM detected significant differences among evolutionary treatments (χ² = 8.701, 2 d.f., *P* = 0.013), as well as significant variation among individual lineages nested within treatments (χ² = 90.512, 8 d.f., *P* < 0.001). Thus, transmission capacity varied both according to the evolutionary environment and among independently evolved lineages.

**FIGURE 4.**
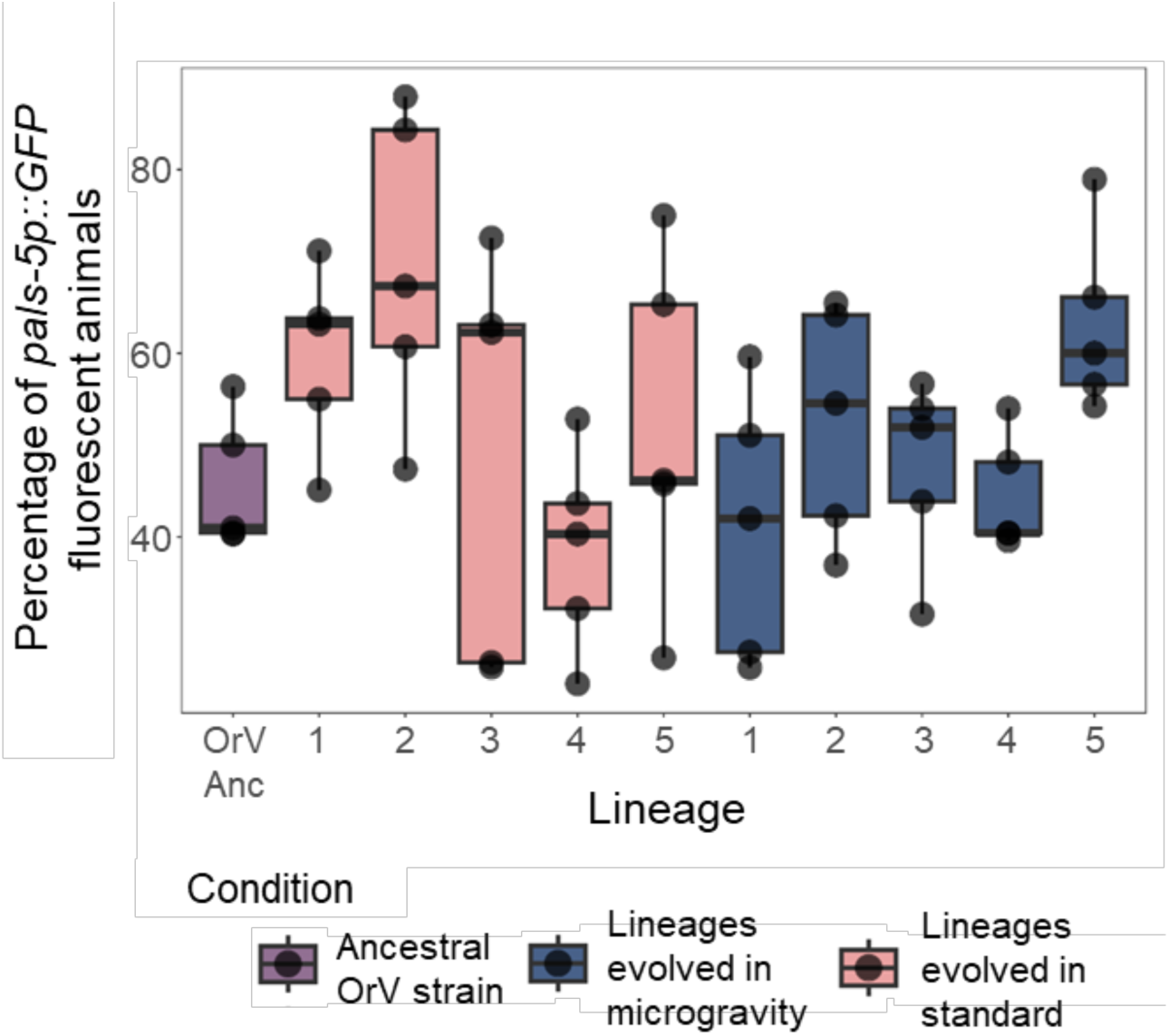
Transmission efficiency measured as the percentage of animals expressing the *pals-5p::GFP* marker after 24 h of exposure to 20 infected animals. The ancestral strain (purple) and the five lineages evolved under standard-gravity (pink) and microgravity (dark blue) conditions were assayed. Transmission experiments were performed with five replicates per lineage.

Pairwise comparisons showed that viruses evolved under standard-gravity transmitted significantly more efficiently than the ancestral strain (0.54 ±0.01 *vs*. 0.45 ±0.03, respectively; Bonferroni-adjusted *P* = 0.030). In contrast, transmission of microgravity-evolved lineages (0.50 ±0.01) did not differ significantly from that of the ancestral virus (Bonferroni-adjusted *P* = 0.192). Therefore, under the standard-gravity assay conditions used here, only the lineages evolved under standard-gravity showed a detectable increase in transmission relative to the ancestor.

This pattern contrasts with the replication results described above. Whereas replication capacity declined in both sets of evolved lineages, transmission increased only after evolution under standard-gravity. This suggests that replication and transmission did not change in parallel during the short-term evolution experiment. However, because transmission was measured only under standard-gravity conditions, the absence of a significant increase in microgravity-evolved lineages should not be interpreted as evidence that these lineages failed to adapt to microgravity. Rather, these results show that microgravity-evolved viruses did not display enhanced transmission when tested outside their selective environment. Reciprocal transmission assays under microgravity would be required to determine whether these lineages acquired environment-specific transmission advantages.

### 3.4. Evolved OrV lineages do not detectably alter host reproductive output under standard-gravity assay conditions

Finally, we evaluated whether evolution under standard-gravity or simulated microgravity conditions altered the effect of OrV infection on host reproductive output. Total progeny production was used as a proxy for host fitness and, therefore, as an inverse measure of virulence. Assays were performed under standard-gravity conditions using animals inoculated with either the ancestral OrV strain or with each evolved viral lineage. Two reference controls were also included: non-inoculated animals and animals inoculated with the ancestral virus. For all viral treatments, inocula were normalized to contain the same number of OrV RNA2 copies.

Across treatments, total progeny production showed only modest differences (Fig. 5). Animals inoculated with the ancestral strain produced, on average, 260 ±87 progeny, whereas non-inoculated animals produced 245 ±88 progeny. Animals infected with lineages evolved under standard-gravity produced 218 ±38 progeny, while those infected with microgravity-evolved lineages produced 226 ±41 progeny. Thus, although evolved lineages tended to show slightly lower mean progeny production than the ancestral strain, variation among replicates was substantial and the differences were small relative to the dispersion of the data.

**FIGURE 5.**
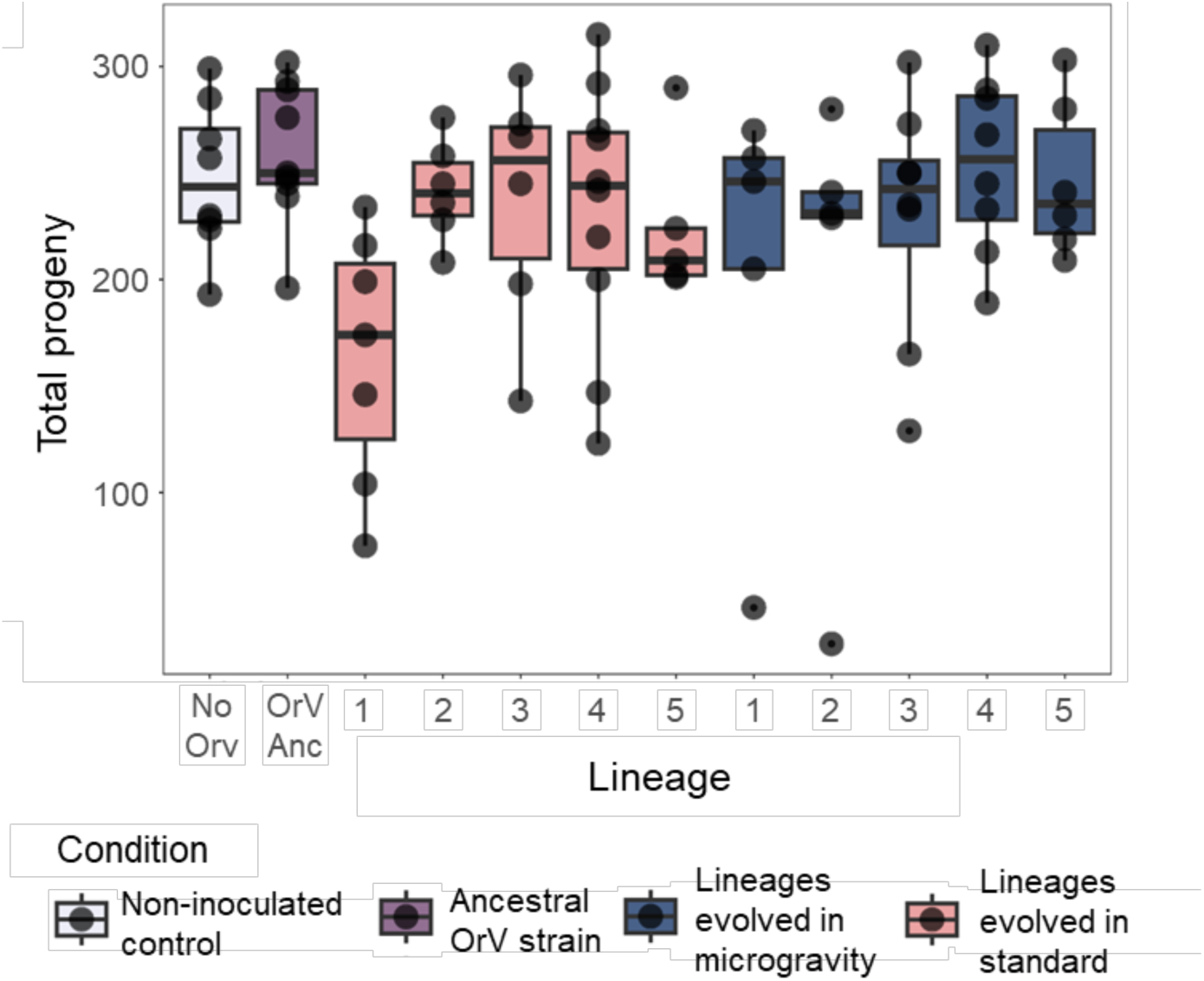
Total progeny across the reproductive lifespan of non-inoculated animals (white), animals inoculated with ancestral OrV (purple), and animals inoculated with the five viral lineages evolved under standard-gravity (pink) or microgravity (dark blue). Animals were inoculated at the L1 stage, and all viral stocks were normalized to equal numbers OrV RNA2 copies. Sample sizes ranged from five to ten biological replicates.

Consistent with this pattern, the GLM detected no statistically significant differences among groups (χ² = 0.281, 3 d.f., *P* = 0.964). Therefore, under the standard-gravity assay conditions used here, neither standard-gravity-nor microgravity-evolved OrV lineages produced a detectable change in host reproductive output relative to the ancestral virus or the non-inoculated control.

These results indicate that the evolutionary changes observed in replication and transmission were not accompanied by measurable changes in host reproductive output. This apparent decoupling suggests that variation in viral replication or transmission does not necessarily translate into detectable differences in this host fitness trait, at least under the inoculum dose, time frame, and assay conditions used here. However, the absence of a detectable effect should be interpreted cautiously. The relatively low and normalized inoculum used in the assay may have reduced the ability to detect subtle differences in virulence, and all measurements were performed under standard-gravity. Consequently, these data do not exclude the possibility that microgravity-evolved lineages differ in their effects on host fitness when assayed under microgravity conditions.

### 3.5 Viral load shows the strongest environment-dependent evolutionary signal among measured traits

To compare the evolutionary signal across the three measured traits, we estimated several quantitative genetic parameters for progeny production, transmission efficiency and viral load in animals infected with OrV lineages evolved under standard-gravity or simulated microgravity conditions. Specifically, we estimated 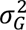, *h*^2^, *S*, *R*, and *n_min_*) These analyses were used to evaluate whether the two evolutionary environments differed in the amount of phenotypic variation available to selection and in the predicted evolutionary response of each trait.

For host reproductive output, the estimated parameters were broadly similar between the two evolutionary treatments. The 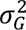 did not differ significantly between standard-gravity and microgravity-evolved lineages, and *h*^2^ showed only a nonsignificant tendency to be higher in the standard-gravity treatment. Likewise, the estimated *S* and predicted *R* did not differ significantly between treatments. The estimated *n_min_* was low in both cases and statistically indistinguishable between environments. These results are consistent with the phenotypic analysis of progeny production, which detected no significant differences among animals infected with ancestral, standard-gravity-evolved, or microgravity-evolved viruses under standard-gravity assay conditions.

Transmission efficiency also showed limited evidence for environment-dependent differences in quantitative genetic structure. Although lineages evolved under standard-gravity showed increased transmission relative to the ancestral virus, the estimated 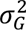, *h*^2^, *S*, *R*, and *n_min_* did not differ significantly between standard-gravity and microgravity-evolved lineages. Thus, despite the phenotypic increase in transmission observed after evolution under standard-gravity, the quantitative genetic estimates did not provide strong evidence that the underlying evolutionary potential for transmission differed between the two evolutionary environments.

In contrast, viral load was the only trait showing clear differences between standard-gravity and microgravity-evolved lineages in several quantitative genetic parameters. The 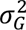 was significantly lower in microgravity-evolved lineages than in standard-gravity-evolved lineages, and *h*^2^ was also significantly reduced under microgravity. The estimated *S* differed significantly between treatments, with a stronger negative value under microgravity. However, the predicted *R* was similar between environments, suggesting that differences in *S* and *h*^2^ partly compensated for one another. In addition, the estimated *n_min_* was significantly higher for viral load in microgravity-evolved lineages, suggesting a more complex genetic or phenotypic basis for variation in viral accumulation after evolution under simulated microgravity.

Overall, these analyses indicate that viral load carries the strongest environment-dependent evolutionary signal among the traits measured. Progeny production and transmission efficiency showed either no significant differences or only limited evidence of treatment-specific quantitative genetic structure. By contrast, viral load differed between evolutionary environments in 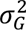, *h*^2^, *S*, and *n_min_*. These results are consistent with the phenotypic assays showing that replication capacity was most strongly affected by the evolutionary treatment. Nevertheless, because these estimates were obtained from traits measured under standard-gravity assay conditions, they should be interpreted as evidence that prior evolution under microgravity altered the expression and structure of viral-load variation in a common environment, rather than as a direct demonstration of adaptation to microgravity.

**Table 1.**
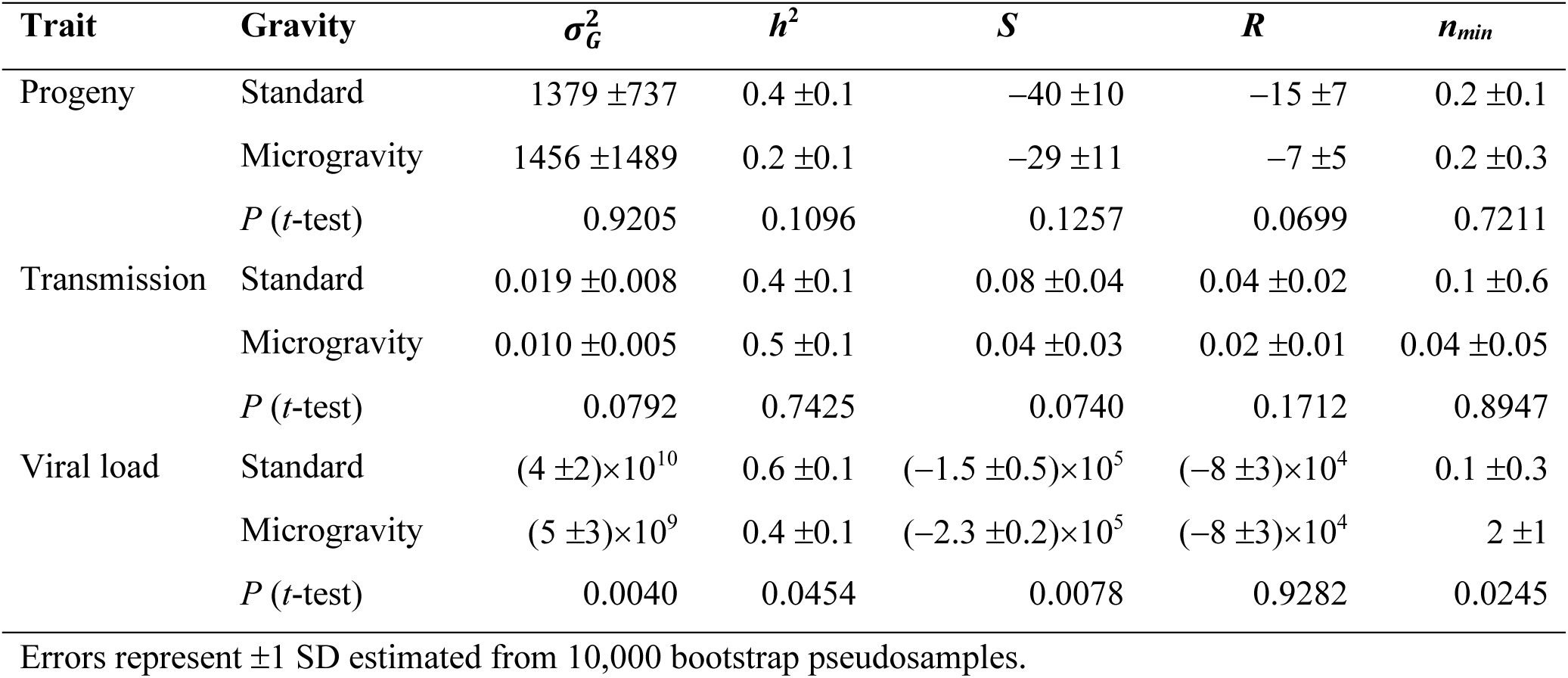
Quantitative genetics analyses for traits measured in animals infected with OrV evolved under the gravity conditions.

| Trait | Gravity | $\sigma_G^2$ | $h^2$ | $S$ | $R$ | $n_{min}$ |
| --- | --- | --- | --- | --- | --- | --- |
| Progeny | Standard | 1379 ± 737 | 0.4 ± 0.1 | -40 ± 10 | -15 ± 7 | 0.2 ± 0.1 |
| | Microgravity | 1456 ± 1489 | 0.2 ± 0.1 | -29 ± 11 | -7 ± 5 | 0.2 ± 0.3 |
| | $P$ ($t$-test) | 0.9205 | 0.1096 | 0.1257 | 0.0699 | 0.7211 |
| Transmission | Standard | 0.019 ± 0.008 | 0.4 ± 0.1 | 0.08 ± 0.04 | 0.04 ± 0.02 | 0.1 ± 0.6 |
| | Microgravity | 0.010 ± 0.005 | 0.5 ± 0.1 | 0.04 ± 0.03 | 0.02 ± 0.01 | 0.04 ± 0.05 |
| | $P$ ($t$-test) | 0.0792 | 0.7425 | 0.0740 | 0.1712 | 0.8947 |
| Viral load | Standard | (4 ± 2) $\times 10^{10}$ | 0.6 ± 0.1 | (-1.5 ± 0.5) $\times 10^5$ | (-8 ± 3) $\times 10^4$ | 0.1 ± 0.3 |
| | Microgravity | (5 ± 3) $\times 10^9$ | 0.4 ± 0.1 | (-2.3 ± 0.2) $\times 10^5$ | (-8 ± 3) $\times 10^4$ | 2 ± 1 |
| | $P$ ($t$-test) | 0.0040 | 0.0454 | 0.0078 | 0.9282 | 0.0245 |

Errors represent $\pm 1$ SD estimated from 10,000 bootstrap pseudosamples.

## 4 Discussion

Here, we investigated whether simulated microgravity alters the short-term evolutionary dynamics of the RNA virus OrV in its natural host, *C. elegans*. To this end, OrV populations were serially passaged for six rounds in hosts maintained either under standard-gravity or simulated microgravity conditions. We then compared evolved lineages for several phenotypic traits associated with viral performance and infection outcome, including replication, transmission, and effects on host reproductive output. Overall, our results indicate that prior evolution under simulated microgravity was associated with altered viral-load dynamics and reduced replication when lineages were subsequently assayed under standard-gravity conditions. However, the magnitude of phenotypic change was generally modest, and the experimental design does not allow a direct test of local adaptation to microgravity.

Given their large population sizes and high mutation rates, RNA viruses typically exist as genetically diverse populations composed of related variants distributed around one or more dominant genotypes. In such systems, multiple variants may coexist in mutation-selection balance, and environmental change can perturb this equilibrium by favoring genotypes that were previously rare under ancestral conditions (Domingo et al. 1996; Arias et al. 2004; Dolan et al. 2018). In this context, simulated microgravity could influence viral evolution either directly, through effects on virion stability, assembly, replication, or transmission, or indirectly, by altering host physiology and thereby modifying the intracellular and organismal environment in which the virus replicates. Because hosts in the microgravity treatment were acclimated for two generations before infection, our design was intended to reduce the contribution of acute host stress responses and to focus on viral evolution in a more stable microgravity-associated host environment. Nevertheless, the present experiment cannot fully distinguish direct effects of microgravity on the virus from indirect effects mediated by host physiology.

Our study differs in several ways from the previous experimental evolution study of OrV performed by Castiglioni et al. (2024). In that work, OrV was evolved for ten passages, with each passage lasting six days. This longer infection window likely provided more opportunities for viral replication and transmission, but also allowed overlapping host generations. In contrast, our experiment used a 44 h infection period, corresponding to larval development of the host, in order to minimize overlapping host generations and reduce the potential for host-virus coevolution during the experiment. This design allowed us to focus on short-term viral evolutionary responses, but it also limited the number of viral replication cycles and transmission opportunities within each passage. In addition, the experiment was restricted to six passages because of limited access to the random positioning machine. These constraints are important for interpreting the results, because limited passage number, short infection duration, and repeated transfers may reduce the opportunity for beneficial mutations to arise and increase in frequency.

Across serial passages, viral load fluctuated markedly in both standard-gravity and microgravity treatments. However, mean viral accumulation was lower under simulated microgravity, a pattern consistent with our previous results showing reduced OrV accumulation during larval development under microgravity conditions (Villena-Giménez et al. 2026a). The temporal dynamics of viral load also showed substantial among-lineage variation, particularly under microgravity. These patterns indicate that simulated microgravity altered the ecological context in which viral populations replicated during experimental evolution. However, lineage-specific temporal trends were generally weak, and average slopes did not differ significantly between treatments. Therefore, these passage-level data are best interpreted as evidence that microgravity modifies short-term viral-load dynamics, rather than as definitive evidence of directional adaptation during the six-passage experiment.

Endpoint phenotypic assays revealed that evolved lineages had lower replication levels than the ancestral virus when tested under standard-gravity conditions. This reduction was observed for both standard-gravity- and microgravity-evolved lineages, with the strongest decrease in the latter. Because inocula were normalized across treatments, these differences likely reflect changes in replication capacity under the assay conditions. The decline in viral load after serial passage may be consistent with the effects of bottlenecks and genetic drift, which can reduce viral fitness during repeated population transfers, particularly in RNA viruses (Chao 1990; Duarte et al. 1992; Lázaro et al. 2003; de la Iglesia and Elena 2007; Zwart and Elena 2015). The stronger decline observed in microgravity-evolved lineages may suggest that the microgravity environment imposed additional constraints on viral amplification, either through changes in host physiology or through direct effects on viral particles or replication processes (Rotem et al. 2018; Macdonald et al. 2024). However, because replication was not assayed under microgravity, we cannot determine whether these lineages would show improved, reduced or unchanged performance in the environment in which they evolved.

Transmission efficiency showed a different pattern from replication. Lineages evolved under standard-gravity conditions transmitted more efficiently than the ancestral virus, whereas microgravity-evolved lineages did not show a statistically significant increase in transmission when tested under standard-gravity. This divergence suggests that replication and transmission did not evolve in parallel during the experiment. In standard-gravity-evolved lineages, reduced replication was accompanied by increased transmission, which may reflect a trade-off between within-host accumulation and between-host spread. Such trade-offs are central to classical models of virulence and transmission evolution (Alizon et al. 2009; Geoghegan and Holmes 2018) and have been documented in several viral systems, including the attenuation and transmission dynamics of myxoma virus in wild rabbits (Kerr 2012). In contrast, microgravity-evolved lineages showed reduced replication without a detectable transmission advantage under standard-gravity assay conditions. One possible explanation is that the microgravity environment modifies host behavior or physiology in ways that alter selection on viral shedding or transmission; for example, microgravity has been shown to affect movement, cytoskeletal organization, and metabolic processes in *C. elegans* (Higashibata et al. 2016). Nevertheless, the absence of increased transmission in microgravity-evolved lineages should not be interpreted as evidence that these viruses failed to adapt to microgravity. Rather, it indicates that these lineages did not display enhanced transmission outside their selective environment. Reciprocal assays under both standard-gravity and microgravity conditions would be required to determine whether transmission evolved in an environment-specific manner.

In contrast to replication and transmission, host reproductive output was not detectably altered by infection with evolved viral lineages. Total progeny production did not differ significantly among non-inoculated animals, animals infected with the ancestral virus, and animals infected with standard-gravity- or microgravity-evolved lineages. Following the definition of virulence as host harm (Geoghegan and Holmes 2018), total progeny provides a useful proxy for host fitness in *C. elegans* and has been widely used in studies of nematode life history and host-pathogen interactions (Barrière and Félix 2005; Villena-Giménez et al. 2026a). The absence of detectable differences suggests that the phenotypic changes observed in viral replication and transmission were not accompanied by measurable changes in this proxy of virulence. Such decoupling between viral accumulation and virulence has been reported in other systems, where viral load does not necessarily predict the magnitude of host fitness effects (Stewart et al. 2005; Cuevas et al. 2015). Several non-mutually exclusive explanations may account for this result. The normalized inoculum used here may have generated relatively mild infections, limiting the ability to detect differences in host reproductive output. In addition, fertility and reproductive output in *C. elegans* can be robust to perturbation because of developmental and genetic buffering (Milloz et al. 2008; Duveau and Félix 2012; Saltzman et al. 2018; Melero et al. 2025). Finally, altered host physiology under stress could modify the relationship between infection and host fitness, as observed in other systems where environmental stress can shift host-virus interactions along the parasitism-mutualism continuum (González et al. 2021). However, because all reproductive assays were performed under standard-gravity conditions, these data do not exclude the possibility that evolved lineages differ in their effects on host fitness when assayed under microgravity.

The quantitative genetic analyses were broadly consistent with the phenotypic results. Progeny production and transmission efficiency showed limited evidence of environment-dependent differences in 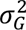, *h*^2^, *S*, predicted *R*, or *n_min_*. In contrast, viral load was the trait with the clearest treatment-associated signal. Microgravity-evolved lineages showed lower 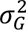and *h*^2^ for viral load, together with a stronger *S* and a higher estimated *n_min_*. These results suggest that prior evolution under simulated microgravity altered the structure of variation underlying viral accumulation when measured in a common standard-gravity environment. The higher estimated *n_min_* may indicate a more complex genetic or phenotypic basis for viral accumulation after evolution under microgravity, consistent with the idea that environmental stress can reshape viral fitness landscapes and the distribution of effects among variants (Dessau et al. 2012; McGee et al. 2016). However, as with the phenotypic assays, these estimates do not directly demonstrate adaptation to microgravity because the relevant traits were not measured under microgravity conditions.

A key limitation of the present study is therefore that all endpoint fitness-associated traits were evaluated exclusively under standard-gravity conditions, irrespective of the environment in which the viral lineages evolved. This design allowed us to compare lineages in a common assay environment, but it precludes a direct test of local adaptation to microgravity and prevents the detection of genotype-by-environment interactions. Consequently, the reduced replication and lack of increased transmission observed in microgravity-evolved lineages under standard-gravity conditions cannot be unambiguously interpreted as evidence of constrained adaptation. These patterns may instead reflect reduced performance outside the selective environment, while potential fitness advantages under microgravity remain untested. Future experiments should therefore include reciprocal assays in which ancestral, standard-gravity-evolved, and microgravity-evolved lineages are phenotyped under both standard-gravity and microgravity conditions.

A second limitation is the short duration and modest evolutionary scope of the experiment. Six passages may be sufficient to detect early phenotypic divergence, but they may not provide enough time for consistent adaptive changes to arise, increase in frequency, and become phenotypically detectable. In addition, repeated population bottlenecks during transfer, short infection periods, and possible limits to the generation or maintenance of genetic diversity could reduce the efficiency of selection. These factors may explain why the magnitude of phenotypic change was generally small and why some traits, particularly host reproductive output and transmission in microgravity-evolved lineages, showed weak or nonsignificant responses. Longer experiments, larger effective population sizes, deeper sequencing of evolved populations, and direct measurements of genetic diversity would help distinguish between genuine evolutionary constraints and insufficient evolutionary opportunity.

Taken together, our results show that simulated microgravity can modify the short-term evolutionary dynamics of OrV, particularly with respect to viral accumulation and replication-associated traits. The strongest and most consistent signal was observed for viral load, whereas transmission and host reproductive output showed weaker or more environment-specific responses. These findings support the use of the *C. elegans*–OrV system as a tractable model for studying how spaceflight-relevant environmental conditions may influence host-virus interactions and viral evolution. At the same time, the current results should be interpreted as evidence of early phenotypic divergence rather than definitive proof of adaptation to microgravity. Determining whether microgravity selects for locally adapted viral phenotypes will require reciprocal fitness assays, longer-term evolution experiments, and genomic analyses of the variants that arise and segregate under altered gravitational environments.

## Data availability statement

The datasets and R code generated during the current study are available in the Zenodo repository, https://doi.org/10.5281/zenodo.20179003.

## Ethics statement

The manuscript presents research on animals that do not require ethical approval for their study.

## Author contributions

A.V-G.: conceptualization, data curation, formal analysis, investigation, methodology, validation, writing-original draft, writing-review and editing; V.G.C.: conceptualization, investigation, methodology, supervision, validation, writing-review and editing; S.F.E.: conceptualization, data curation, formal analysis, funding acquisition, project administration, supervision, writing-original draft, writing-review and editing. The funders played no role in study design, data collection, analysis and interpretation of data, or the writing of this manuscript.

## Acknowledgements

We thank Francisca de la Iglesia (I^2^SysBio) and Rebecca Hernández-Antolín (LSC) for excellent technical support. We are deeply grateful to Carlos P. Garay (LSC) for his commitment and dedication to research in subterranean biology, as well as his constant material and emotional support.

## Generative AI statement

During manuscript revision, generative AI tools were used to assist with language editing, readability, and organization of selected sections. The authors critically reviewed, edited, and approved all AI-assisted text and take full responsibility for the final content of the manuscript.

---

**Supplementary Table S1.** Fitting of viral load time-series data to the ARIMA(1,0,0) model.

| Lineage | Cox $\lambda$ | $\phi$ | SE $\phi$ | $C$ | SE $C$ | BIC | $Q^*$ | $P$ | slope | 95% CI | | Kendall $\tau$ | $P$ |
| --- | --- | --- | --- | --- | --- | --- | --- | --- | --- | --- | --- | --- | --- |
|  |  |  |  |  |  |  |  |  |  | lower | upper |  |  |
| Standard 1 | 1.983 | 0.452 | 0.203 | 17.179 | 3.356 | 111.66 | 10.204 | 0.017 | 0.011 | −0.044 | 0.114 | 0.033 | 0.880 |
| Standard 2 | 2.000 | 0.515 | 0.194 | 14.151 | 2.694 | 100.60 | 6.861 | 0.077 | 0.066 | 0.003 | 0.126 | 0.386 | 0.028 |
| Standard 3 | −0.900 | 0.644 | 0.178 | 0.901 | 0.012 | −102.41 | 0.808 | 0.848 | −0.011 | −0.037 | 0.035 | −0.163 | 0.363 |
| Standard 4 | −0.900 | 0.602 | 0.186 | 0.890 | 0.012 | −100.11 | 1.395 | 0.707 | 0.052 | −0.026 | 0.073 | 0.199 | 0.289 |
| Standard 5 | 2.000 | 0.569 | 0.201 | 16.216 | 4.186 | 112.71 | 5.947 | 0.114 | 0.002 | −0.080 | 0.096 | 0.020 | 0.940 |
| Microgravity 1 | 2.000 | 0.976 | 0.029 | 1.966 | 0.996 | 91.74 | 5.589 | 0.133 | −0.013 | −0.065 | 0.098 | −0.020 | 0.940 |
| Microgravity 2 | −0.900 | 0.581 | 0.185 | 0.862 | 0.011 | −101.76 | 3.509 | 0.320 | 0.054 | −0.003 | 0.077 | 0.333 | 0.058 |
| Microgravity 3 | 2.000 | 0.660 | 0.187 | 14.720 | 3.221 | 95.64 | 2.030 | 0.566 | −0.016 | −0.055 | 0.060 | −0.190 | 0.289 |
| Microgravity 4 | 2.000 | 0.508 | 0.202 | 16.455 | 1.610 | 81.16 | 7.500 | 0.058 | 0.067 | 0.022 | 0.083 | 0.451 | 0.010 |
| Microgravity 5 | 2.000 | 0.478 | 0.202 | 20.800 | 3.753 | 113.85 | 7.046 | 0.071 | −0.028 | −0.078 | 0.026 | −0.242 | 0.173 |

$Q^*$ : Ljung-Box tests with 3 d.f.

